# Lack of phenotypic variation despite population structure in larval utilization of pea aphids by populations of the lady beetle *Hippodamia convergens*

**DOI:** 10.1101/740506

**Authors:** Christy Grenier, Bryce Summerhays, Ryan Cartmill, Tanairi Martinez, Roxane Saisho, Alexander Rothenberg, Alicia Tovar, Andrew Rynerson, Jerrika Scott, John J Obrycki, Arun Sethuraman

**Affiliations:** Department of Biological Sciences, California State University San Marcos, San Marcos, California; Department of Biology, Benedict College, Columbia, South Carolina; Department of Entomology, University of Kentucky, Lexington, Kentucky

## Abstract

The convergent lady beetle (*Hippodamia convergens*) is a generalist natural enemy that is utilized extensively in augmentative biological control across the United States. Recent studies have pointed to both genetic and phenotypic differences in Western (California) versus Eastern (Kansas) populations of the species. Here we investigate (1) genetic population structure, and (2) phenotypic differences in the utilization of pea aphids at temperatures that resemble the Western United States in (a) Eastern versus Western populations, (b) F1 Eastern X Western hybrids versus their progenitor populations, and investigate the effects of interaction between (c) Eastern and Western populations. We found no differences in final pupal weight, or the net weight gain ratio through larval development from the third instar to pupal stage, despite genetic population structure. Our study points towards plastic response and effectiveness in feeding phenotypes of Eastern and Western populations of *H. convergens*, and the absence of hybrid vigor and heterozygote advantages in hybrids.

## Introduction

Ladybird beetles (also known as ladybugs, Coleoptera: Coccinellidae) are commonly utilized as natural enemies against infestation of aphids, whiteflies, and scales across the world (Roy and Wajnberg 2008). In North America, the convergent lady beetle, *Hippodamia convergens* is the most abundant native species of coccinellids used in both introduction and augmentative biological control (Bjørnson 2008). Western populations disperse into the Sierra Nevada Mountains to form large overwintering aggregations (Wheeler and Ring 2014). These large concentrations of adults make the Western population easily susceptible to unregulated collections, which are then sold to farmers or home gardeners and released across the United States (Obrycki and Kring 1998; Sethuraman et al., 2015).

Recent population genetic studies of *H. convergens* across their range in the continental United States have revealed the presence of at least two structured geographic populations (termed Western and Eastern populations - Sethuraman et al., 2015). *H. convergens* within their designated Western and Eastern populations in the United States have likely adapted to varying natural conditions, including pathogens and parasitoid cycles. These populations have also previously been shown to have differences in developmental histories, overwintering behavior, and reproductive diapause (Hagen 1962, Obrycki and Tauber 1982, Obrycki et al., 2001). Research from Obrycki and Tauber (1982) show that unlike Western populations of *H. convergens*, Eastern populations develop slower during warmer periods in early spring, but faster later in spring. Despite their differences, Eastern and Western populations are able to hybridize with each other without any known reproductive barriers (Obrycki et al., 2001). Many coccinellid beetles are multivoltine, producing two or more broods within a year which would allow these beetles to mate before migrating back to their respective sites (Koch and Hutchison 2003). Augmenting populations by bringing Western and Eastern populations together to create hybrids can also potentially increase the fitness of the hybrid population, a phenomenon that is commonly described as ‘hybrid vigor’ (Seko et al., 2012). However, no comparative studies of the utilization of aphids by Western, Eastern, or hybrid populations of *H. convergens* under native or nonnative climates have been conducted. This type of study is needed to quantify the potential levels of aphid biological control resulting from augmentative releases of the Western populations of *H. convergens*.

Biological control, while providing effective control of agricultural pests, comes at the cost of, or is affected by several non-target effects. Modern agricultural practices often utilize pesticides against crop pests, but have adverse consequences for the pests and other beneficial arthropods which are utilized in biological control (Biondi et al., 2012a, 2012b, Geiger et al., 2010), and contribute to global arthropod decline (Potts et al., 2010, Benton et al., 2002, Hallmann et al., 2017). Transportation and augmentation of *H. convergens* populations has also led to the movement and spread of arthropod pathogens and parasitoids (Bjørnson 2008). Studies have been conducted in California to document the effects of native augmentative releases of *H. convergens* (Flint et al., 1995; Flint and Dreistadt, 2005). However, little is known about the effectiveness of transporting Western collected populations of *H. convergens* throughout the United States.

The objective of this study is to understand the effectiveness of human mediated augmentation of predatory *H. convergens* from the Western population on the Eastern population, and how potential interaction between Eastern and Western populations might differentially affect levels of biological control. With the differences between populations, do environmental and development factors affect how they interact with each other through their predation of pea aphids? Does hybridization between the two inbred populations increase the ability of removing pests in agricultural use due to hybrid vigor? Using both Eastern and Western populations of *H. convergens* lady beetles found in the United States, as well as F1 Eastern x Western hybrids, we address the following questions: (1) Do inbred Eastern and Western populations differ in their effectiveness of utilization of pea aphids?, (2) Are F1 Eastern x Western hybrids more effective at the utilization of pea aphids than their progenitor populations?, and (3) Is there an effect from interaction between Eastern and Western populations? Previous studies have shown that the adult body weight of *H. convergens* beetles are positively correlated with fecundity and the number of aphids consumed during larval development (Kajita and Evans 2010; Obrycki et al., 2001). Thus we address these questions by (1) assessing the pupal weight and weight gain by use of a net weight gain ratio (Final weight – Initial Weight/Initial Weight) of genetically disjunct Western and Eastern larvae when placed individually on aphid bearing plants (2) raising F1 Eastern x Western hybrid larvae to assess their pupal weight and net weight gain ratio when individually placed on an aphid bearing plant, compared to the pupal weight of the Western and Eastern populations under the same conditions, and (3) assessing the pupal weight and net weight gain ratio of Western, and Eastern beetles when one beetle from both populations was placed on the same plant. Additionally, we ascertain genotypic differences between Western and Eastern populations of *H. convergens* using microsatellite genotyping and analyses of population structure.

## Methods

*H. convergens* were raised from field collected beetle egg masses from Kansas (provided by JP Michaud, Kansas State University), representing the Eastern Population of the species. The Western population of *H. convergens* were field collected from adult aggregations on Palomar Mountain in San Diego County in Southern California. Beetles were raised on frozen or live pea aphids (*Acyrthosiphon pisum*), which were reared on fava bean plants (*Vicia faba*), in a greenhouse at California State University San Marcos, San Marcos, CA. The greenhouse temperatures had an average high temperature of 27.7 °C and average low temperature of 16.3 °C from January to May 2019. Western and Eastern populations were started from approximately 45 individuals, and were inbred for several generations before beginning experimental crosses. At least 5 Eastern virgin females were crossed with Eastern males, and at least 5 Western virgin females were crossed with Western males for the within population crosses. At least 4 virgin females were crossed with the opposing population to make F1 Eastern x Western hybrids. Mating pairs were allowed 48 hours to mate, after which the males were separated, and females were fed pea aphids *ad libitum*, and allowed 48 hours to lay egg masses. Once the egg masses were laid, females were removed, and egg masses were collected in preparation for the experimental assays.

To assess for competition and biocontrol efficacy, a common-garden setup was utilized. A fava bean plant (~10 cm in height, 2 week old sapling, with 7±1 leaves) was placed in a 2 liter plastic bottle with a cut-out black mesh window. Third instars from the crossing experiments were separated into individual cups, and starved for 24 hours prior to the beginning of our assay. After 24hrs, 0.050 ± 0.003g of aphids (approximately 50 aphids) were placed inside each bottle and allowed approximately 3-12 hours to settle and infect the fava bean plant. Thereon, third instar larvae of similar weight (average difference for all pairs was 0.000225g, average initial weight for all individuals was 0.005653g) were weighed using an analytical balance and then placed inside the following treatment bottles: 1) 1 Western larva, 2) 1 Eastern larva, 3) 1 F1 Western X Eastern hybrid larva, 4) 2 Western larvae, 5) 2 Eastern larvae, 6) 2 F1 Eastern x Western hybrid larvae, and 7) 1 Western larvae with 1 Eastern Larvae which were painted with acrylic paint to determine the individuals. Treatments 1-3 were used to assess phenotypic differences between Western, Eastern, and F1 Western x Eastern hybrids without interaction. Treatments 4-6 were used as a control to see that there was no difference when two individuals of the same population interacted versus no interaction in treatments 1-3. Treatment 7 was used to assess phenotypic differences when Western and Eastern larvae interacted with one another.

Larvae were then weighed every other day with an analytical balance until pupation, where the weight of the pupa would be the final weight recorded (approximately 8 days). On the fourth day, fava bean plants inside the bottles were watered, and another approximate 0.050g of pea aphids were placed inside each bottle to ensure each fava bean plant still had aphids, and that all larvae had *ad libitum* access to food. The experiments were repeated until eight sets of replicates were completed from March-May 2019. Results from treatments where larvae had gone missing or died were eliminated from statistical analyses.

### Population Genetic Structure

To assess the population structure of Western and Eastern populations that were used in this study, we performed genotyping at six polymorphic microsatellite loci *sensu* Sethuraman et al., 2015. 5 Western and 6 Eastern adult beetles from the study were flash frozen with liquid nitrogen and whole genomic DNA was extracted using Qiagen DNeasy Kits using the manufacturer’s protocol. The six microsatellite loci used in this study were developed and characterized previously by Sethuraman et al., 2015 (Table 3). PCR’s were performed using the KAPA Taq ReadyMix PCR Kit (Kit code KK1006) in a final total volume of 25μL containing 144.2 ± 31.5ng of genomic DNA, 1X KAPA Taq ReadyMix at 1.5 mM MgCl2, and 0.3μM of each primer (fluorescently labeled using 6-FAM dye set on the 5’ end of the forward primer). PCR reaction conditions were as follows: 95°C for 3 min followed by 35 cycles at 95°C for 30 s, 30 s at primer specific annealing temperatures (see Table 3), and 72°C for 20 s and a final extension period at 72°C for 20 s. PCR products were then visualized on a 2% agarose gel to ensure quality of bands. Samples with high quality amplicons were then genotyped via capillary electrophoresis at Retrogen (San Diego, CA) using the Life Technologies’ DS-33 dye set and GS600LIZ size standard for sizing fragments of length 20-600 bp.

### Microsatellite Data Analysis

All raw fragment files were analyzed using ABI PeakScanner v.1.0 and genotypes were ascertained by three independent reviewers, to minimize bias. These genotypes were then analyzed for population structure using two methods: (1) using only the 11 individuals from this study, and (2) using all individuals from California, Kansas from the study of Sethuraman et al., 2015, and the 11 individuals from this study. Genotypes were converted into the GENIND format and analyzed for population structure using DAPC (Jombart et al., 2010) using the R package adegenet (Jombart and Ahmed 2011). The number of presumed subpopulations or clusters (commonly denoted by K) was varied from K = 1 to 10, and the optimal number of subpopulations explained by the data were assessed using the Bayesian Information Criterion (BIC) reported by the adegenet package (see Fig. 3(A), and 4(A)). Population structure was then visualized using stacked bar plots of admixture proportions (here denoted as “membership probability”).

### Statistical Analyses

Performance of Eastern,Western, and hybrid populations of *H. convergens* was assessed using the beetles’ final pupation weight, and weight gain as a ratio (Final weight – Initial Weight/Initial Weight) as a proxy for fitness. A mean net weight gain was calculated, when it was not possible to distinguish individuals in conspecific larval replicates. All statistical analyses were performed using R (version 3.6.3). For each test, if the data was not normal, a log or square root transformation used. One-way ANOVAs were performed (Table 1, and 2) to test the following hypotheses: 1) The Western population of *H. convergens* is better at pea aphid utilization than the Eastern under climate conditions that mimic the Western United States, 2) Due to hybrid vigor, the F1 Eastern x Western hybrid population will be better at utilizing pea aphids than both the Eastern and Western population, and 3) in the interaction assay, a Western *H. convergens* larvae will show greater pupal weight and net weight gain compared to an Eastern *H. convergens* larva.

**Table 1.**
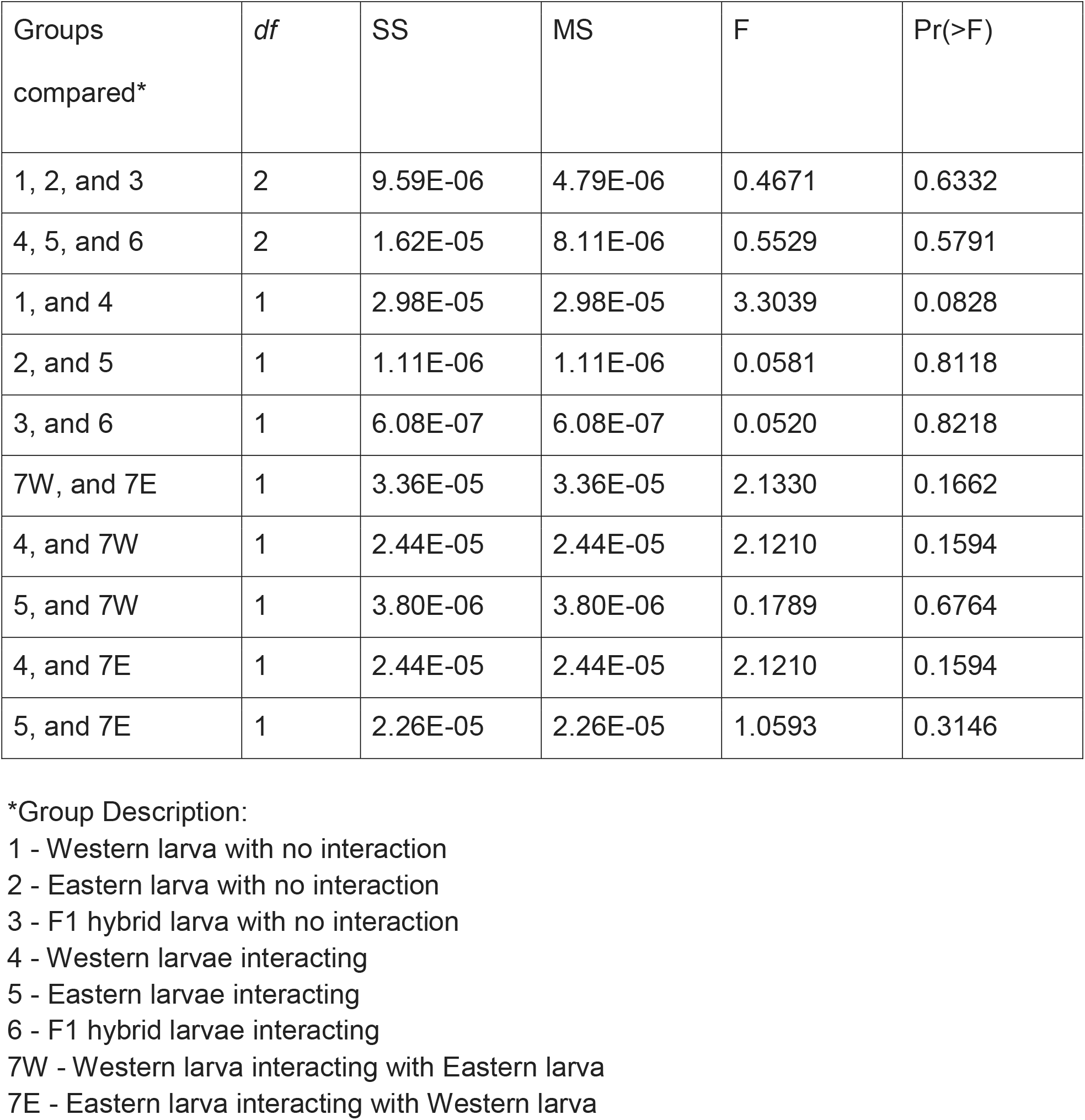
One-way ANOVA results when comparing final pupal weight of single larvae, and interacting larvae, of Western, Eastern, and F1 Western X Eastern hybrid larvae of *H. convergens*. For all statistical tests pupal weight was used, and Shapiro-Wilks test indicated normal distribution. All statistical tests were performed using R (version 3.6.3).

**Table 2.**
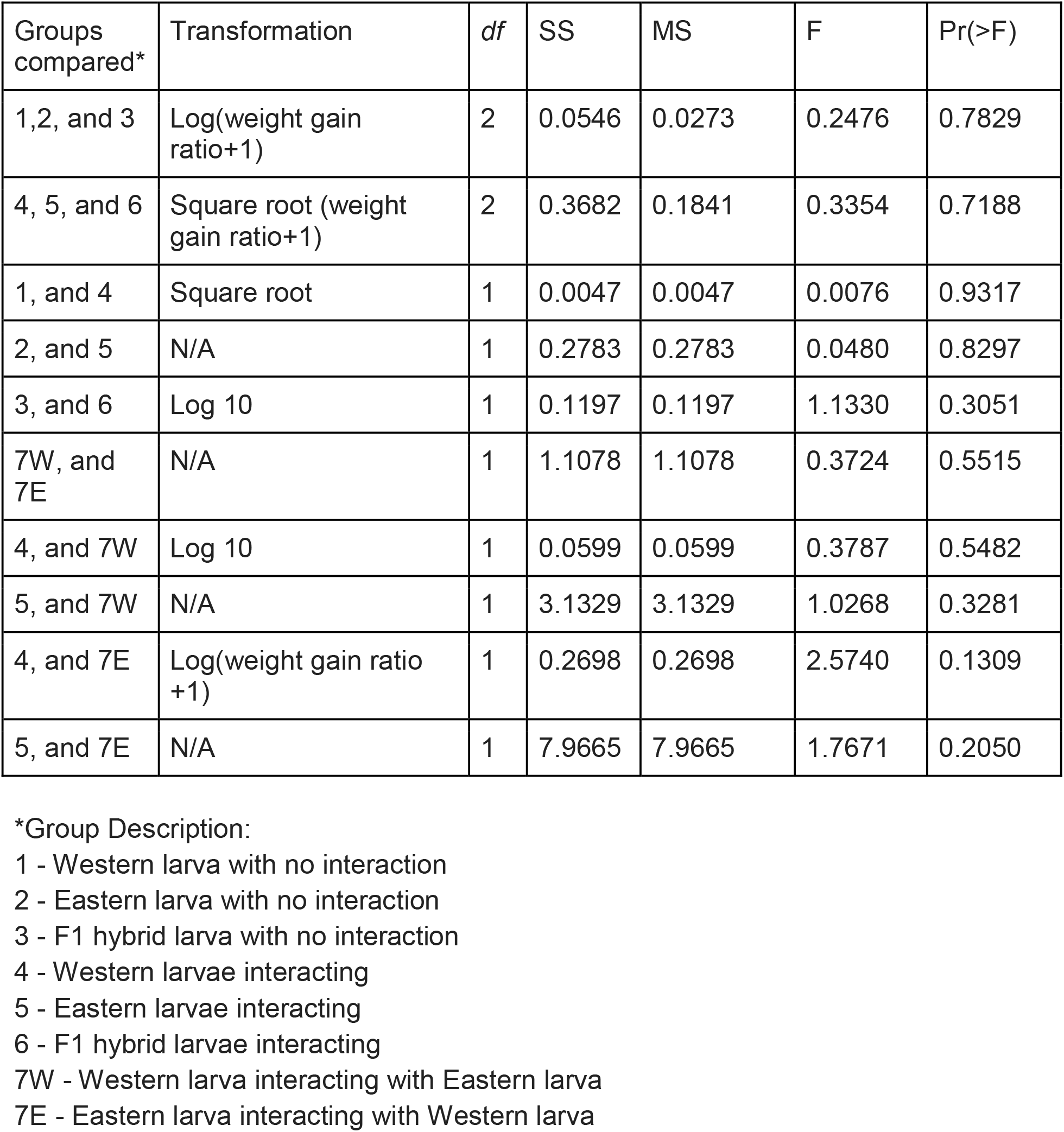
One-way ANOVA results when comparing the net weight gain ratio of lone larvae, and interacting larvae, of Western, Eastern, and F1 Western X Eastern hybrid larvae of *H. convergens*. Weight gain ratio (Final weight – Initial Weight/Initial Weight) was used for all statistical tests, and data were transformed and normalized when applicable for every statistical test. All ANOVAs and Shapiro-Wilks tests were performed using R (version 3.6.3).

| Groups compared* | Transformation | df | SS | MS | F | Pr(>F) |
| --- | --- | --- | --- | --- | --- | --- |
| 1,2, and 3 | Log(weight gain ratio+1) | 2 | 0.0546 | 0.0273 | 0.2476 | 0.7829 |
| 4, 5, and 6 | Square root (weight gain ratio+1) | 2 | 0.3682 | 0.1841 | 0.3354 | 0.7188 |
| 1, and 4 | Square root | 1 | 0.0047 | 0.0047 | 0.0076 | 0.9317 |
| 2, and 5 | N/A | 1 | 0.2783 | 0.2783 | 0.0480 | 0.8297 |
| 3, and 6 | Log 10 | 1 | 0.1197 | 0.1197 | 1.1330 | 0.3051 |
| 7W, and 7E | N/A | 1 | 1.1078 | 1.1078 | 0.3724 | 0.5515 |
| 4, and 7W | Log 10 | 1 | 0.0599 | 0.0599 | 0.3787 | 0.5482 |
| 5, and 7W | N/A | 1 | 3.1329 | 3.1329 | 1.0268 | 0.3281 |
| 4, and 7E | Log(weight gain ratio +1) | 1 | 0.2698 | 0.2698 | 2.5740 | 0.1309 |
| 5, and 7E | N/A | 1 | 7.9665 | 7.9665 | 1.7671 | 0.2050 |

\*Group Description:
1 - Western larva with no interaction
2 - Eastern larva with no interaction
3 - F1 hybrid larva with no interaction
4 - Western larvae interacting
5 - Eastern larvae interacting
6 - F1 hybrid larvae interacting
7W - Western larva interacting with Eastern larva
7E - Eastern larva interacting with Western larva

## Results

### Population Structure

Two loci (Hcv13 and Hcv30) failed chi-squared tests of Hardy-Weinberg Equilibrium (p < 0.05, Table 3). Nonetheless, since analyses of population structure using DAPC are agnostic to deviations from HWE, these were included in further analyses. Estimates of population genetic structure using microsatellite markers revealed the separation of our field-sampled Eastern (Kansas) and Western (California) populations into K = 3 subpopulations, when analyzed by themselves (Fig. 3A). Importantly, the Western population was classified as a unique subpopulation, compared to the Eastern population, which was further split into 2 subpopulations (Fig. 3B). However, analysis of population structure using our field-sampled beetles in combination with Californian and Kansan beetles from the study of Sethuraman et al., 2015 inferred the presence of K = 2 subpopulations (Fig. 4A). In these analyses, our Eastern and Western populations clustered together with some Californian and Kansan populations from the study of Sethuraman et al., 2015 (Fig. 4B).

**Table 3.**
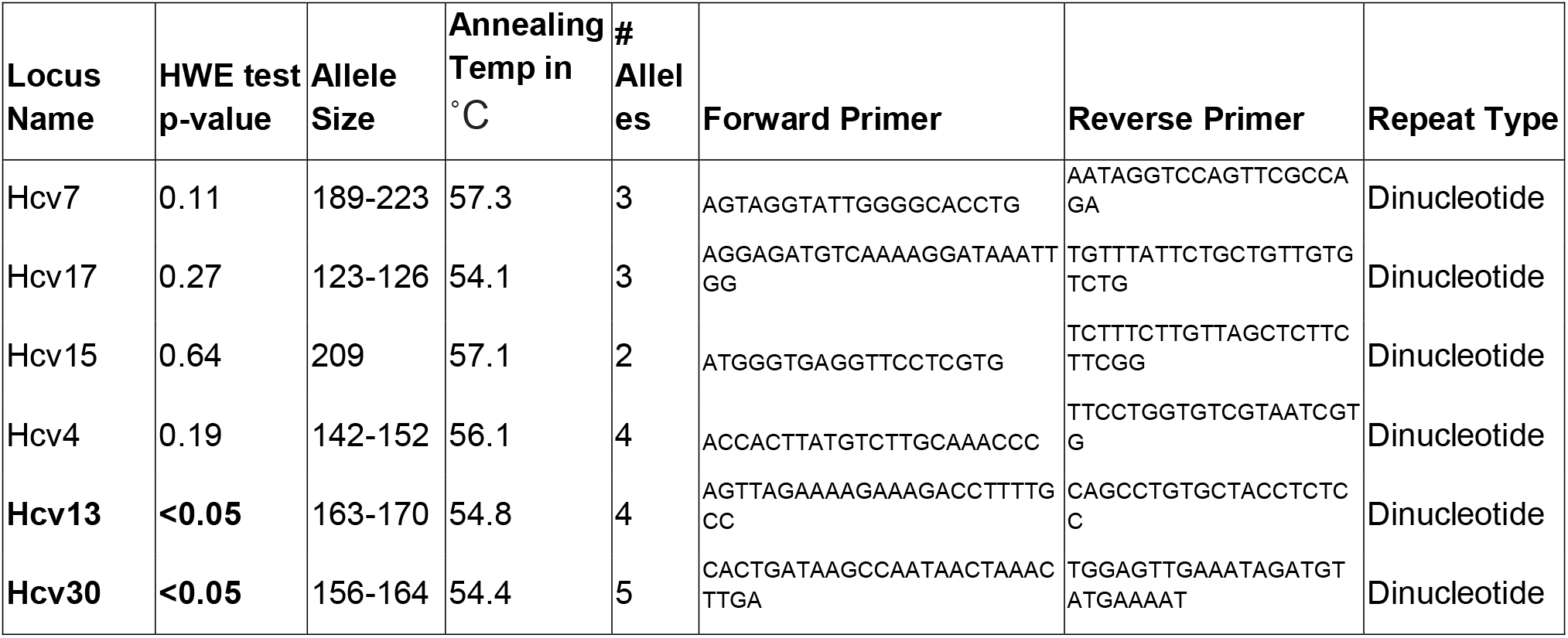
List of primers, allele sizes, and population genetic summary statistics of each locus used in the microsatellite analyses to deduce population structure of Eastern and Western populations of *H. convergens*. Loci that fail the chi-squared test of HWE are shown in boldface.

### Interaction Experiments

In all treatments, *H. convergens* larvae molted successfully and pupated into adults. Data was transformed if needed and Shapiro-Wilks tests indicated no deviation from normality (P>0.05) for all data sets used for ANOVA. Outliers did not have a significant effect on the results, and were not excluded from the statistical tests (Fig. 1, and 2). The difference between pupal weight of the larvae when individually placed on pea aphid bearing plants was not statistically significant between the Western, Eastern, and F1 hybrid populations (d.f.=2, F=0.4671, and P=0.6332, Table 1), nor was it statistically significant for the net weight gain (d.f=2, F=0.2474, and P=0.7829, Table 2). There was no statistical difference between pupal weight of a single individual vs pupal weight of interacting individuals from the same population in Western, Eastern, or F1 hybrids (d.f.=1, F=3.3039, P=0.0828, d.f=1, F=0.0581, P=0.8118, and d.f.=1, F=0.0520, P=0.8218 respectively, Table 1), and the net weight gain showed no difference as well (d.f.=1, F=0.0076, P=0.9317, d.f.=1, F=0.0480, P=0.8297, and d.f.=1, F=1.1330, and P=0.3015 respectively, Table 2). Furthermore, both pupal weight, and the net weight gain ratio of within-population interactions showed no difference between Western, Eastern, or F1 hybrids (P>0.5). Comparison of pupal weight between an Eastern individual interacting with a Western individual when placed on the same plant also bore no statistical significance for pupal weight or the net weight gain (d.f.= 1, F= 0.2133, P=0.1662, and d.f.=1, F=0.3724, P=0.5515 respectively, Table 1 and 2).

**Fig.1.**
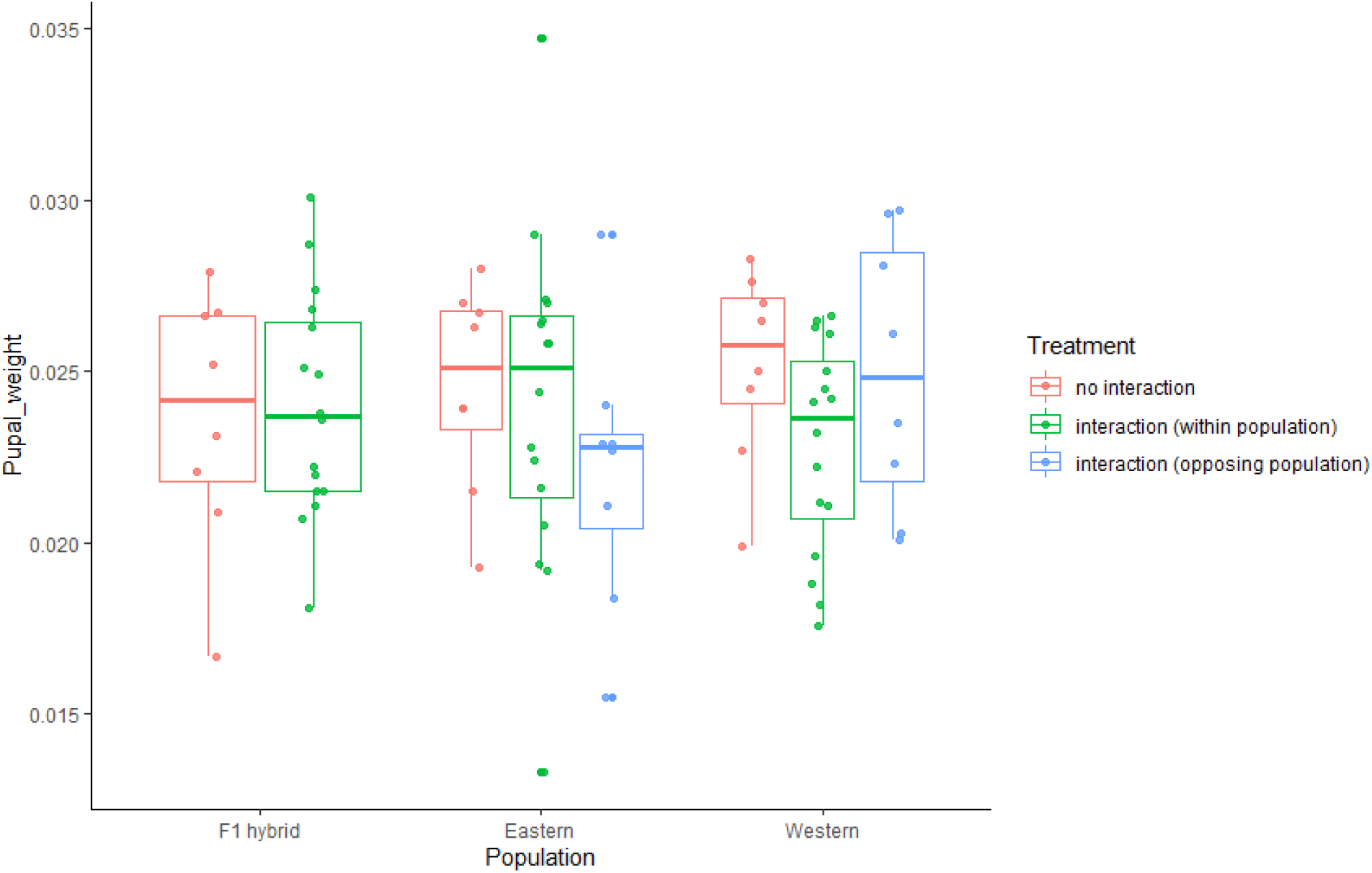
Distribution of pupal weights of larvae with no interaction, and interaction of F1 hybrids, Eastern, and Western larvae.

**Fig.2.**
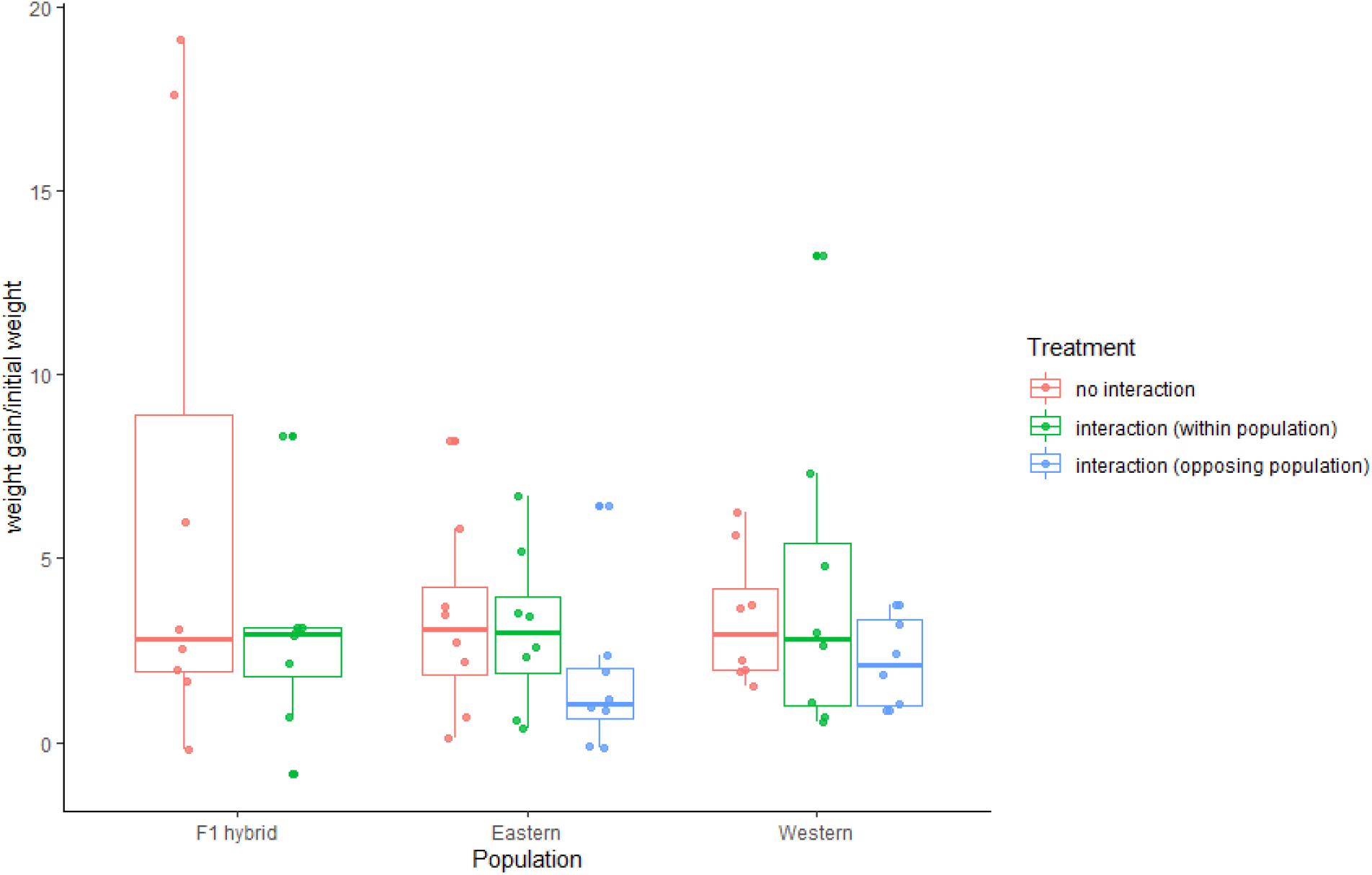
The distribution of net weight gain ratio of larvae with no interaction, and interaction of F1 hybrids, Eastern, and Western larvae.

## Discussion

Several species of terrestrial arthropods are utilized extensively as natural enemies against common agricultural pests and invasive species, and are estimated to result in billions of dollars in agricultural savings across the globe (Coombs et al 1996, Huang et al., 2018). Within the United States, beneficial insects provide over 50 billion dollars worth of services and 4.5 billion is attributed to biological control organisms known as natural enemies (Landis et al., 2008). Natural enemies can have lasting effects by establishing populations in the introduced ranges for long term recurring utilization of crop pests (Enkerli et al., 2004). Oftentimes, biological control involves introducing non-native species into a new range (Dodd, Alan 1959), which interact with a diverse array of intra- and inter-specific competitors (Evans 1991). Quantifying their effectiveness is therefore of great importance to biological control programs (Tauber and Tauber 1975, Evans 1991).

Within the United States, it is common practice to transfer populations of Western *H. convergens* to the Eastern United States to control aphid pest infestations. This augmentative biological control can lead to hybridization of Eastern and Western populations (Sethuraman et al., 2015). When genetically structured populations hybridize the hybrid progeny are known to exhibit physical traits and behavioral phenotypes that confer greater fitness than their progenitor populations (Seko et al., 2012, Li et al., 2018). Both Eastern and Western populations are known to migrate over large geographical distances, however it is unknown if mating occurs between the populations during migration, and if the populations return to their previous locations (Sethuraman et al., 2015). This was further elucidated by analyzing the population genetics of our Eastern and Western populations. Our analyses clearly separate our Eastern and Western populations of *H. convergens* into unique clusters (Fig. 3B). Despite our Eastern and Western populations being genetically structured into separate populations when compared to each other, we did not find a significant difference in aphid utilization between F1 hybrids nor Eastern or Western populations (P>0.78). However, inclusion of a previous microsatellite dataset from Sethuraman et al., 2015 cannot conclusively rule out the absence of hybridization between our Eastern and Western populations owing to them structuring together with other previously analyzed Californian, and Kansan populations (Fig. 4B).

**Fig.3.**
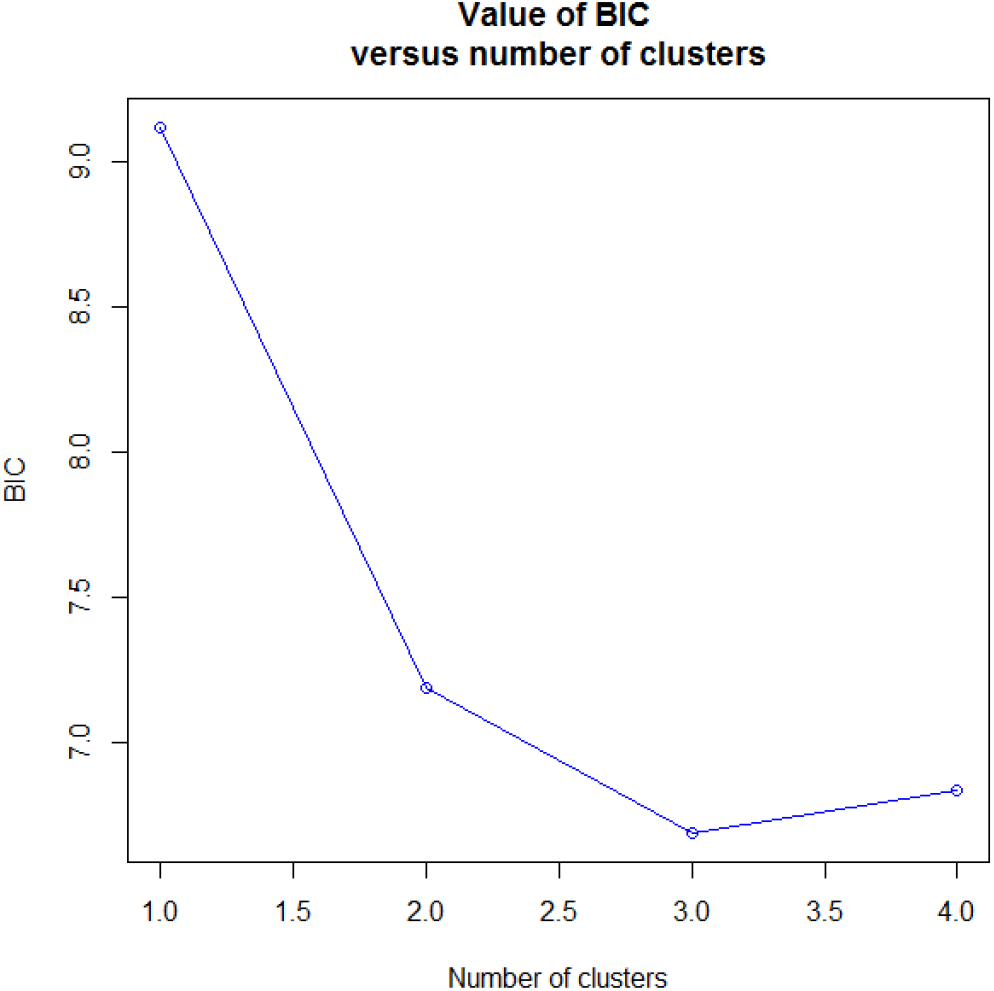
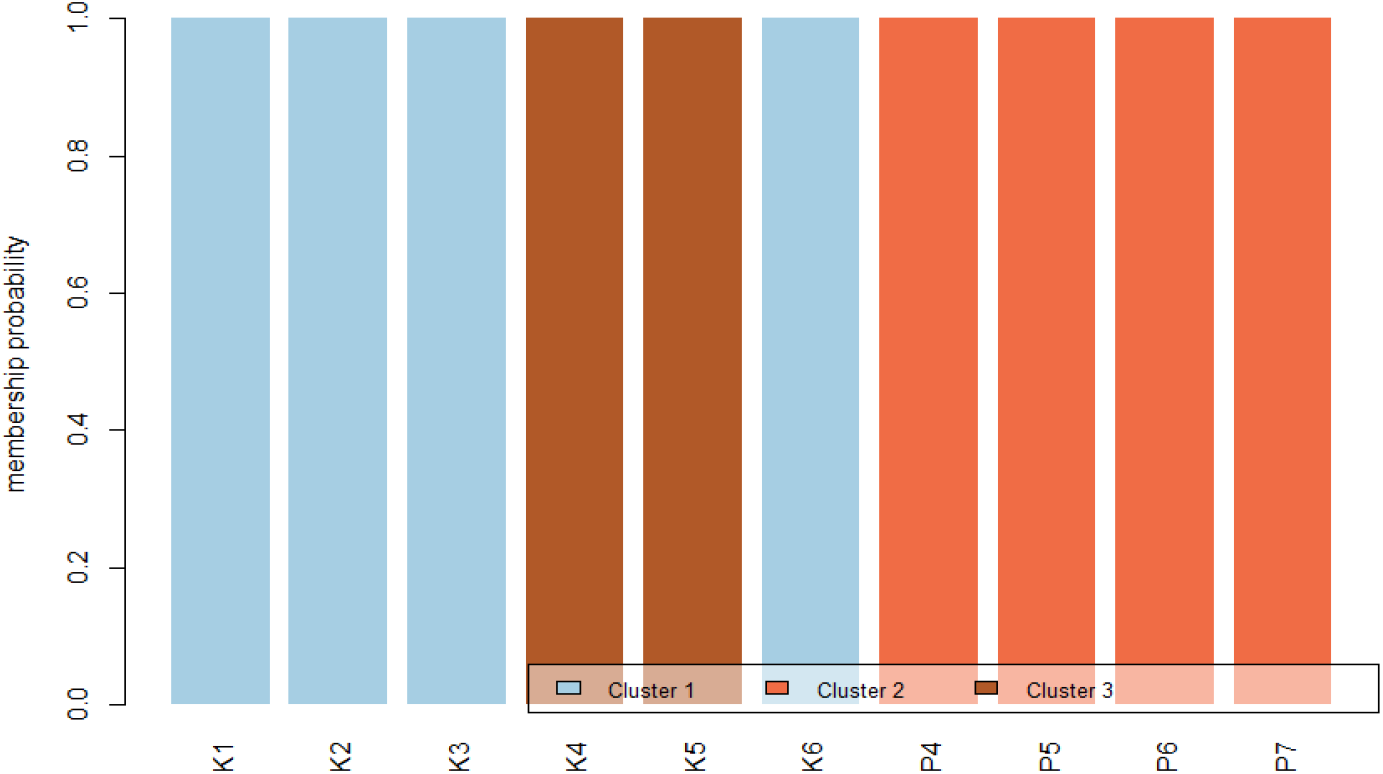
**(A)** Bayesian Information Criterion (BIC) for clustering Eastern and Western populations from this study into one of K = 4 populations. **(B)** Membership probabilities (admixture proportions) estimated for our Eastern beetles (K1-K6), and Western beetles (P4-P7) to one of K=3 populations. All Western beetles cluster together into one subpopulation, whereas the Eastern beetles are split into two separate subpopulations.

**Fig.3.**
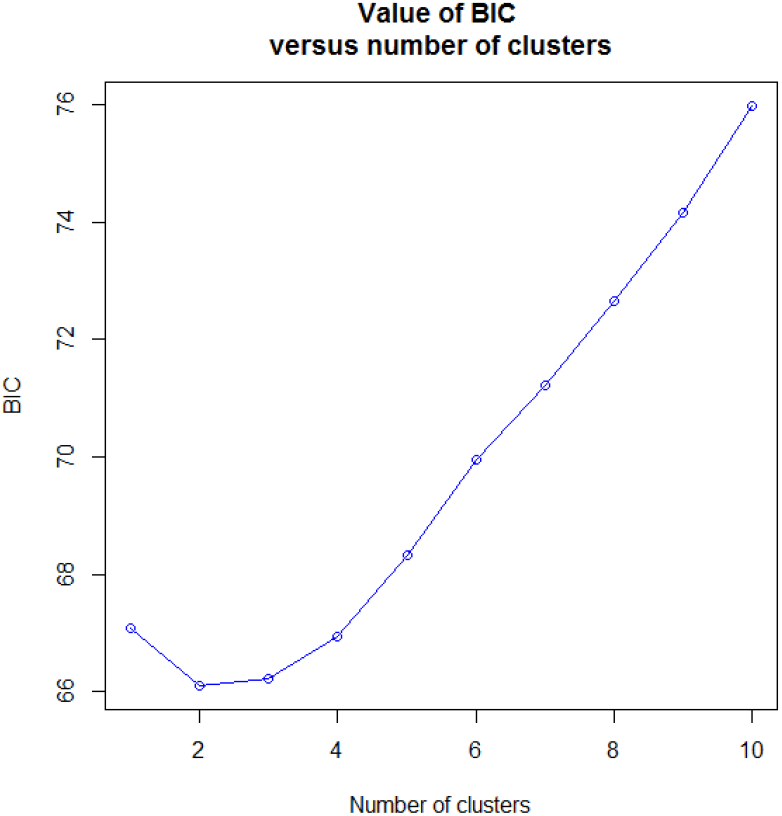
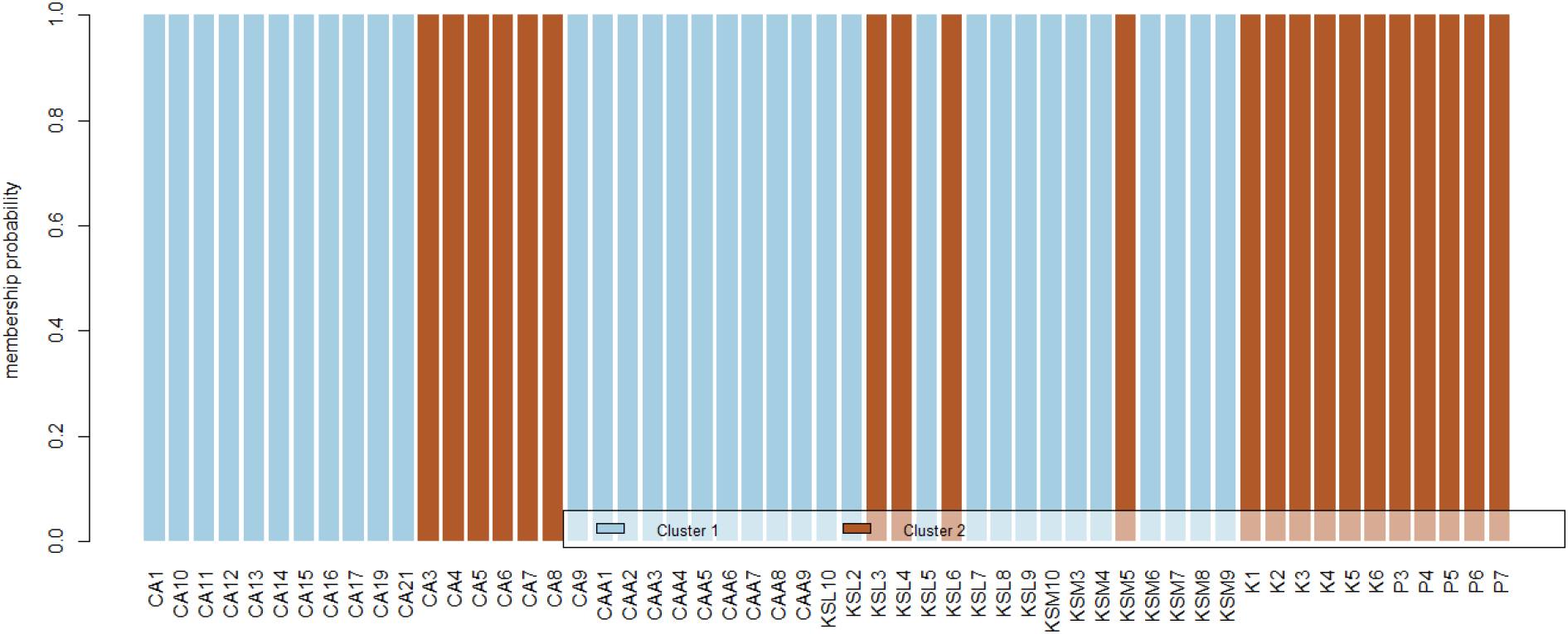
**(A)** Bayesian Information Criterion (BIC) for clustering Eastern and Western populations from this study into one of K = 10 populations together with populations from California and Kansas from the study of Sethuraman et al., 2015. **(B)** Membership probabilities (admixture proportions) estimated for our Eastern beetles (K1-K6), and Western beetles (P4-P7) to one of K=2 populations with other California (CA) and Kansas populations (KS). All Western and Eastern populations structure together with beetles from California and Kansas, indicating some degree of hybridization in these populations due to human-mediated augmentation.

Our findings show that the significant phenotypic differences between Eastern and Western populations of *H. convergens* in the United States (Obrycki et al., 2001), become irrelevant in the warmer western conditions, suggesting that importation of beetles to the West, from the East would not lower effectiveness as a biocontrol agent. Evidence of differences in photoperiodic responses have been observed in beetle populations that have not adapted to their local environment, which can result in slower developmental cycles when compared to populations that are native to the area (Obrycki et al., 2018). This response to photoperiods has also been shown to be heritable, indicating that augmentation and importation may also affect the ability for future generations of the introduced population to compete with native populations (Reznik et al., 2017). Similarly, differences in temperature regimes in newer environments could also lead to a difference in reproductive diapause between populations, that can cause introduced populations to develop at a slower rate than the native population (Wang et al., 2013).

Although our results indicate that no disadvantageous or advantageous effects in the control of pea aphids may occur when larvae interact with one another when provided with access to excess aphids; when beetle larvae were starved for 24 hours together, the larger and older instars were found to feed on younger instars. Intraspecies/guild predation is well documented in lady beetles, especially when there is a large size difference between larvae and adults on larva or eggs (Bayoumy et al., 2015, Agarwala and Dixon 1992). We paired most of the larvae to be in their third instar stage, and were approximately similar in size and weight, although 7/32 pairs had a difference in weight of more than double the weight of the smaller individual. However, some larvae escaped, or disappeared from the tent, putatively indicating intraspecific predation. This data was subsequently removed from the study so as to not bias our statistical analyses.

In summary, there were no significant differences between our Eastern, Western, or F1 hybrid populations of beetles, as individuals, or paired, in their effectiveness of utilization of the aphid crop pests (Table 1 and 2). These experiments were conducted in a greenhouse with semi-regulated temperatures in Southern California, an environment which does not mirror the environment of the Eastern Region of the United States. The similarities between Eastern and Western individuals could hence be attributed to testing at higher, Western temperatures in Southern California. Further studies should thus measure the rates of utilization of aphids at lower temperatures that mimic the Eastern Region of the United States. These studies would allow a better understanding of the environmental effects on introduced, or augmented populations of *H. convergens* in the colder versus warmer regions of the United States. Additionally, future experiments addressing interaction between Eastern and Western *H. convergens* would provide useful information in determining the effects of human augmentation of Western lady beetles to Eastern populations.

## Acknowledgments

This work was funded by USDA grant #2017-06423 to Drs. George Vourlitis and Arun Sethuraman. AS was supported by NSF ABI Development grant #1564659. We would like to acknowledge our collaborator, Dr. JP Michaud for field collecting and shipping the Kansas (Eastern) population of *H. convergens*. RS, RC were funded by a CSUSM Summer Scholars fellowship, and JS was funded by an NSF REU grant #1852189 to Drs. Betsy Read and Arun Sethuraman. We would also like to thank Dr. Elinne Becket for help with optimizing the microsatellite analyses.

## Author Contributions

JJO, AS, BS conceptualized the experiments, BS, CG, TM, and RC performed all the experiments, and RS, AR, and JS performed all the statistical analyses and analyses of microsatellite data. AT, AR also performed the microsatellite analyses. Primary author is CG and BS with contributions from RS, AR, JS, RC,TM, AS and JJO.

